# The evolutionary constraints on angiosperm chloroplast adaptation

**DOI:** 10.1101/2022.07.12.499704

**Authors:** Elizabeth Hannah Joan Robbins, Steven Kelly

**Affiliations:** Department of Plant Sciences, University of Oxford, South Parks Road, Oxford, OX1 3RB, United Kingdom

**Keywords:** Chloroplast, Evolvability, Adaptive Evolution, Evolutionary Constraint, Phylogenomics

## Abstract

The chloroplast (plastid) arose via endosymbiosis of a photosynthetic cyanobacterium by a non-photosynthetic eukaryotic cell approximately 1.5 billion years ago. Although the plastid underwent rapid evolution by genome reduction, its rate of molecular evolution is low and its genome organisation is highly conserved. Here, we investigate the factors that have constrained the rate of molecular evolution of protein coding genes in the plastid genome. Through analysis of 773 angiosperm plastid genomes we show that there is substantial variation in the rate of molecular evolution between genes. We show that variation in the strength of purifying selection between genes is a major determinant of variation in the rate of molecular evolution. We further demonstrate that the distance of a gene from the likely origin of replication influences the rate at which it has evolved, consistent with time and distance dependent mutation gradients. In addition, we show that the amino acid composition of a gene product constraints its substitution tolerance, limiting its rate of molecular evolution. Finally, we demonstrate that the mRNA abundance of a gene is a key factor in determining its rate of molecular evolution, suggesting an interaction between transcription and DNA repair in the plastid. Collectively, we show that the location, composition, and expression of a plastid gene can account for ≥32% of the variation in its rate of molecular evolution. Thus, these three factors have exerted a substantial limitation on the capacity for adaptive evolution of plastid genes, and constrained the evolvability of the chloroplast.

## Introduction

The chloroplast (plastid) is a membrane bound organelle that arose from the endosymbiosis of a photosynthetic cyanobacterium by a non-photosynthetic eukaryotic cell approximately 1.5 billion years ago (Schwartz and Dayhoff 1978; Martin and Kowallik 1999; Hedges, et al. 2004). Early in the transition from cyanobacterium to organelle, the plastid underwent a substantial genome reduction whereby thousands of genes were either lost or transferred to the nucleus (Blanchard and Schmidt 1995; Martin, et al. 2002; Kelly 2021). As a consequence, modern-day plastid genomes in plants contain fewer than 5% of the genes found in their free-living bacterial relatives, with the majority of plant plastids containing only ∼80 protein-coding genes (Palmer 1985; Timmis, et al. 2004). As a result, despite comprising over 80% of the protein (Makino and Osmond 1991; Li, et al. 2017) and 80% of the mRNA (Forsythe, et al. 2022) in a plant cell, the combined forces of gene loss and endosymbiotic gene transfer mean that plastids typically contain fewer than 0.5% of the genes found in a typical plant cell.

Although the plastid underwent rapid evolution by genome reduction, the rate of molecular evolution of the plastid encoded genes and their organisation in the plastid chromosome has been markedly slow and highly conserved. Comparisons between plant species has revealed that the plastid genome has a mutation rate that is an order of magnitude lower than that of the nuclear genome (Wolfe, et al. 1987; Smith 2015). The finding that the mutation rate of the plastid was low in comparison to the nuclear genome was initially unexpected given that the plastid genome is predominantly uniparentally inherited (Birky 1995; Mogensen 1996), where an absence of sexual recombination is thought to lead to an accumulation of deleterious mutations in a phenomenon known as Muller’s ratchet (Muller 1964). Accordingly, it was proposed that gene conversion in the presence of a high genome copy number per plastid (∼900 per plastid (Bendich 1987)) acts to counteract the effect of Muller’s ratchet to prevent accumulation of mutations (Birky and Walsh 1992; Khakhlova and Bock 2006). However, it is unclear whether other factors intrinsic to the plastid genome itself, also act to limit the rate of molecular evolution of plastid genes.

One factor that could differentially impact the rate of molecular evolution of plastid encoded genes is their position in the chromosome relative to the point of replication initiation. Most plant species contain a plastid genome that has a quadripartite structure (Palmer 1985; Daniell, et al. 2016) composed of two single-copy regions (the large single-copy region and the small single-copy region) separated by two inverted repeat regions. Although multiple replicative mechanisms have been hypothesised (Kolodner and Tewari 1975; Takeda, et al. 1992; Kunnimalaiyaan and Nielsen 1997; Oldenburg and Bendich 2004; Oldenburg and Bendich 2016), the origin(s) of replication have been repeatedly proposed to be located in the inverted repeat regions, and both the replication mechanism and position relative to these origins are thought to impact the evolution of the nucleotide sequence along the plastid genome (Wolfe, et al. 1987; Smith 2015). For example, adenine to guanine deamination gradients disseminate away from the inverted repeat regions, and mutation rates in the single-copy regions are much higher than in the inverted repeats (Wolfe, et al. 1987; Krishnan and Rao 2009). The formation of the deamination gradients is thought to be attributable to the differential length of time in which the regions of the plastid genome are single-stranded during DNA replication, and the difference in mutation rate between the single-copy and inverted repeat regions is thought to be due to enhanced frequency of homologous recombination mediated repair in the inverted repeat (Khakhlova and Bock 2006). Other context and location dependent effects on mutation in the plastid genome have been described which also alter the rate and type of mutations that occur along the plastid chromosome (Wolfe, et al. 1992; Kelchner 2000; Jansen, et al. 2007; Zhu, et al. 2016; Niu, et al. 2017). However, it is unknown whether these position-dependent effects have influenced the rate of molecular evolution of plastid gene sequences, or how these phenomena interact with other factors to modulate the rate of molecular evolution of plastid encoded genes.

Here, we present a holistic analysis of the molecular evolution of the single-copy plastid encoded protein-coding genes from 773 species of angiosperms. We show that there is substantial variation in the rate of molecular evolution between genes, with genes whose products function in energy production having evolved more slowly than those whose products function in information processing (transcription and translation). We demonstrate that differences in strength of purifying selection acting on plastid encoded genes can explain ∼50% of variation in the rate of molecular evolution between genes. We subsequently perform a holistic analysis of multiple factors that could also contribute to the variation in the rate of molecular evolution between plastid encoded genes, including gene position, amino acid composition, nucleotide composition, levels of gene expression, transcript biosynthetic cost and transcript translational efficiency. In doing so, we reveal that the level of mRNA abundance of a gene, its position relative to the inverted repeat region, and its amino acid composition each significantly constrain the rate of evolution of plastid encoded genes. Thus, in addition to selection to preserve molecular function, the combined action of compositional factors and production requirements impose a severe constraint on the molecular adaptation of the chloroplast.

## Results

### There is a large variation in rate of molecular evolution between plastid located genes

To understand the factors that have influenced the rate of molecular evolution of plastid encoded genes, a dataset of fully sequenced plastid genomes was compiled. To do this, the full set of sequenced plastid genomes was downloaded from NCBI (Wolfsberg, et al. 2001) and subjected to filtration to remove genomes that did not contain a full set of the genes typically found in angiosperms. This resulted in a dataset comprising complete plastid genomes from 773 species of angiosperms, each containing 69 single-copy protein-coding genes that are ubiquitously present in all species. Gene trees were inferred separately from nucleotide and protein multiple sequence alignments for all ubiquitously conserved genes (see methods) and the phylogenetic trees were used to evaluate and compare the rates of molecular evolution for each gene at both the nucleotide and amino acid level, respectively.

The total length of each gene tree was used as the estimate for the rate of molecular evolution for each gene. This measure accounts for differences in alignment length between genes because the branch lengths in the tree are estimates of the number of substitutions per aligned sequence site (Supplementary Figure 1). This measure is also not affected by species sampling, variation in model of sequence evolution, or variation in topology of the underlying tree, as each of these factors was held constant in all analyses that were conducted. Specifically, all 773 plant species are in each gene tree and each gene is inferred from a strictly single-copy gene in each species. Furthermore, the topology of each gene tree was constrained to the consensus topology inferred from a concatenation of all genes (see methods), and the models of sequence evolution inferred from these concatenated alignments were held constant across all individual tree inferences using parameters that allow for comparison of branch lengths between trees. Thus, this analysis removes any potential phylogenetic bias between the rates that are measured. For ease of interpretation, the rate of molecular evolution of the nucleotide sequences and the rate of molecular evolution of the protein sequences are henceforth referred to as the rate of nucleotide evolution (*k*_N_) and rate of protein evolution (*k*_P_), respectively.

Comparison of the rates of molecular evolution between genes at the nucleotide level revealed that there was an 8-fold difference between the fastest and slowest evolving genes, *rpl22* and *psbL*, respectively (Figure 1A, Supplemental Table S1). In contrast, the rate of protein evolution varied much more substantially with a 76-fold difference between the fastest and slowest evolving proteins, *rpl22* and *atpH*, respectively (Figure 1B, Supplemental Table S1). Thus, even though the plastid genome has evolved slowly in comparison to the nuclear genome (Wolfe, et al. 1987; Smith 2015), there is still substantial variation in the rate of molecular evolution between plastid encoded genes.

**Figure 1.**
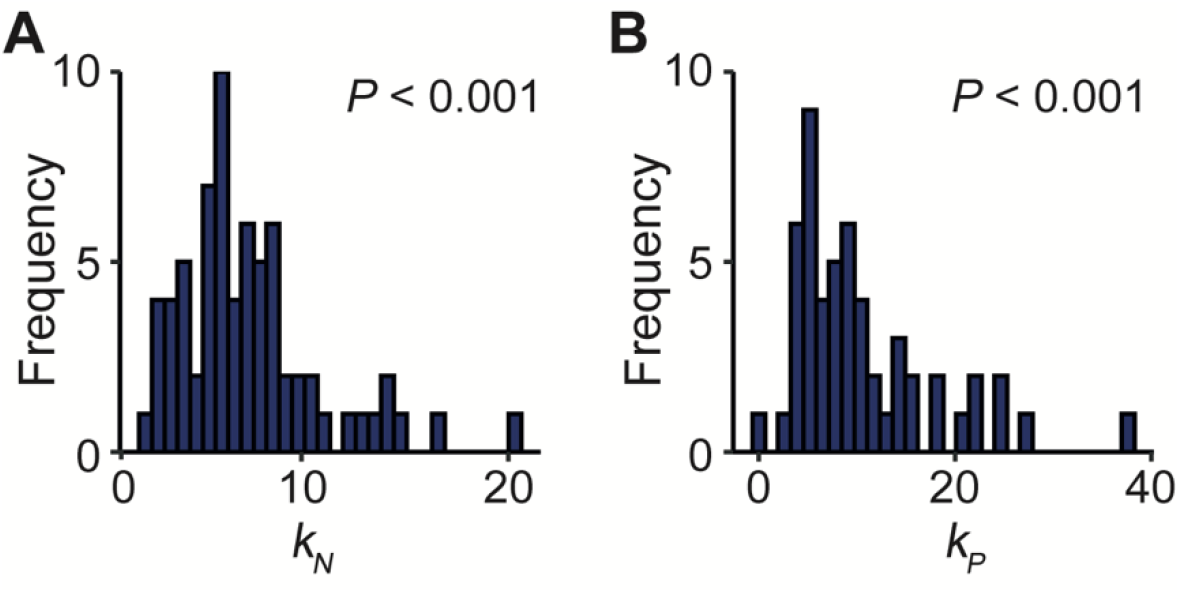
The rates of molecular evolution for 69 single-copy protein-coding plastid genes. **A-B**) Frequency distributions of the rates of nucleotide evolution (*k*_N_) and protein evolution (*k*_P_), respectively. Rates of molecular evolution have the units of the total substitutions per sequence site per tree (S_st_). *P*-values correspond to the result of a Shapiro-Wilk test for normality (all *P* < 0.05).

### Plastid genes with functions in energy production have evolved slower and are more constrained than genes that function in information processing

Inspection of the rank order of the rates molecular evolution (Figure 2A and B) suggested that genes involved in energy production (i.e., genes whose products function in photosystem I, II, cytochrome b_6_f complex, ATP-synthase and the NAD(P)H dehydrogenase-like complex or CO_2_ fixation) had a lower rate of molecular evolution than those involved in information processing (i.e., genes that function in transcription, mRNA maturation, translation or protein degradation). Comparison of these two groups revealed that genes with functions in energy production evolved slower than those with functions in information processing, with mean fold differences of 1.7 and 2.8 for the rate of nucleotide and protein evolution, respectively (Welch’s t-test, *P* < 0.001) (Figure 2C and D). Within each category there was little to no difference in the evolutionary rate of genes between multiprotein complexes (Supplementary Figure 2). Thus, plastid encoded genes with functions in energy production have evolved more slowly than plastid encoded genes with functions in information processing.

**Figure 2.**
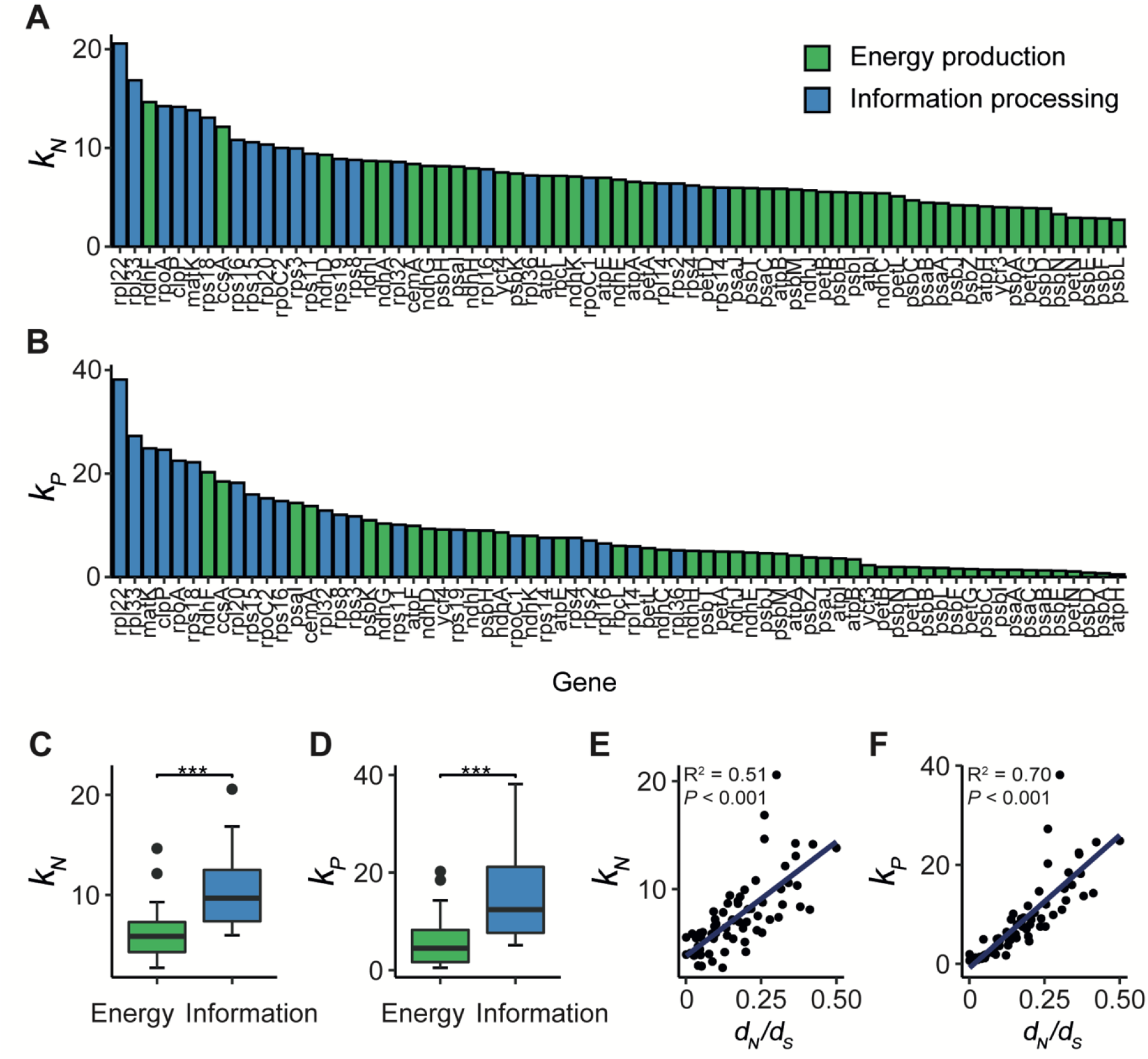
Relationship between gene function and rate of molecular evolution. **A-B**) Bar graphs showing the rate of nucleotide evolution (**A**) and protein evolution (**B**) for individual genes in order of descending value from left to right. Genes are coloured by their high-level function: green, energy production; blue, information processing. The energy production category includes genes that function in, or are associated with photosystem I, photosystem II, cytochrome b_6_f complex, ATP-synthase, NAD(P)H dehydrogenase-like complex or rubisco. The information processing category includes genes involved in mRNA maturation in addition to genes that encode subunits of the 70S ribosome, the plastid encoded RNA polymerase and the plastid protease system. **C-D**) The difference in the rate of nucleotide evolution and protein evolution between genes involved in energy production (green, n=47) versus information processing (red, n=22), respectively. Statistical significance was assessed using a Welch’s t-test (all *P* < 0.001, indicated by ***). **E-F**) Scatter plots of the non-synonymous to synonymous mutation rate (*d*_N_/*d*_S_) versus the rate of nucleotide and protein evolution, respectively. Linear models are shown in blue with their associated R^2^ and *P*-values.

### The strength of purifying selection modulates the rate of plastid gene evolution

It has been previously shown that many genes within the energy production category are subject to strong purifying selection (Zhang, et al. 2020; Sheikh-Assadi, et al. 2022; Ślesak, et al. 2022). Since purifying selection acts to preserve the existing sequence due to functional constraints, it follows on that genes exposed to strong purifying selection are likely to have lower rates of molecular evolution than genes that experience weaker purifying selection. To determine whether this was evident in the dataset presented here, the strength of purifying selection was calculated from the ratio of the rate of non-synonymous mutation to synonymous mutation (*d*_N_/*d*_S_). All genes had a *d*_N_/*d*_S_ ≤ 0.5 indicating strong purifying selection and functional constraint (Kimura 1989), and consistent with previous observations. Although all genes have been subject to strong purifying selection, there was a 500-fold difference observed between the least and most functionally constrained genes (*matK, d*_N_ /*d*_S_ = 0.5; *psbA* and *petB, d*_N_/*d*_S_ = 0.001). As expected, genes under stronger purifying selection had lower rates of molecular evolution and *vice versa*. Variation in the strength of purifying selection could explain 51% of the variance in the rate of nucleotide evolution between plastid encoded genes (R^2^= 0.51, *P* < 0.001) (Figure 2E) and 70% of the variance in the rate of protein evolution (R^2^ = 0.70, *P* < 0.001) (Figure 2F). Thus, variation in the strength of purifying selection between genes is a major component of rate variation between genes, but does not fully account for all variation in the rate of molecular evolution.

### Physicochemical constraints on amino acid substitution have limited the rate of molecular evolution of plastid encoded genes

Given that selection acting to preserve the existing sequence was a major component of variation in the rate of molecular evolution, we sought to determine whether compositional differences between the sequences further exacerbated this constraint. Such a compositional effect may arise from the biochemical differences and similarities between amino acid monomers. Specifically, amino acids are not all equally dissimilar to each other, and many share similarities in their size, charge, and reactivity of their side chains. The extent of similarity between amino acids varies such that each different amino acid monomer has a different tolerance to substitution in protein sequences (Sneath 1966; Epstein 1967; Grantham 1974; Henikoff and Henikoff 1992). This substitution tolerance can be effectively described using substitution frequency data summarised in the BLOSUM62 amino acid substitution matrix (Henikoff and Henikoff 1992). We hypothesised that proteins containing a higher proportion of amino acids that were less tolerant to substitution would have lower rates of molecular evolution because mutations would be more likely to be deleterious to the encoded protein sequence. To investigate this, a substitution tolerance score was calculated for each gene (see methods) and compared to the rate of molecular evolution of that gene. This revealed variation in substitution tolerance between genes could explain significant components of variation in the rate of nucleotide evolution (R^2^ = 0.16, *P* < 0.001) (Figure 3A) and the rate of protein evolution (R^2^ = 0.23, *P* < 0.001) (Figure 3B). Here, genes with lower substitution tolerance scores had lower rates of molecular evolution. The observation that substitution tolerance imposes a stronger constraint on the rate of molecular evolution of the protein sequences than the nucleotide sequences is likely to be due to redundancy in the genetic code permitting mutation in the nucleotide sequence without altering the identity of the encoded amino acid. Thus, the rate at which a plastid gene has evolved during the radiation of the angiosperms is dependent on the amino acid composition of that gene.

**Figure 3.**
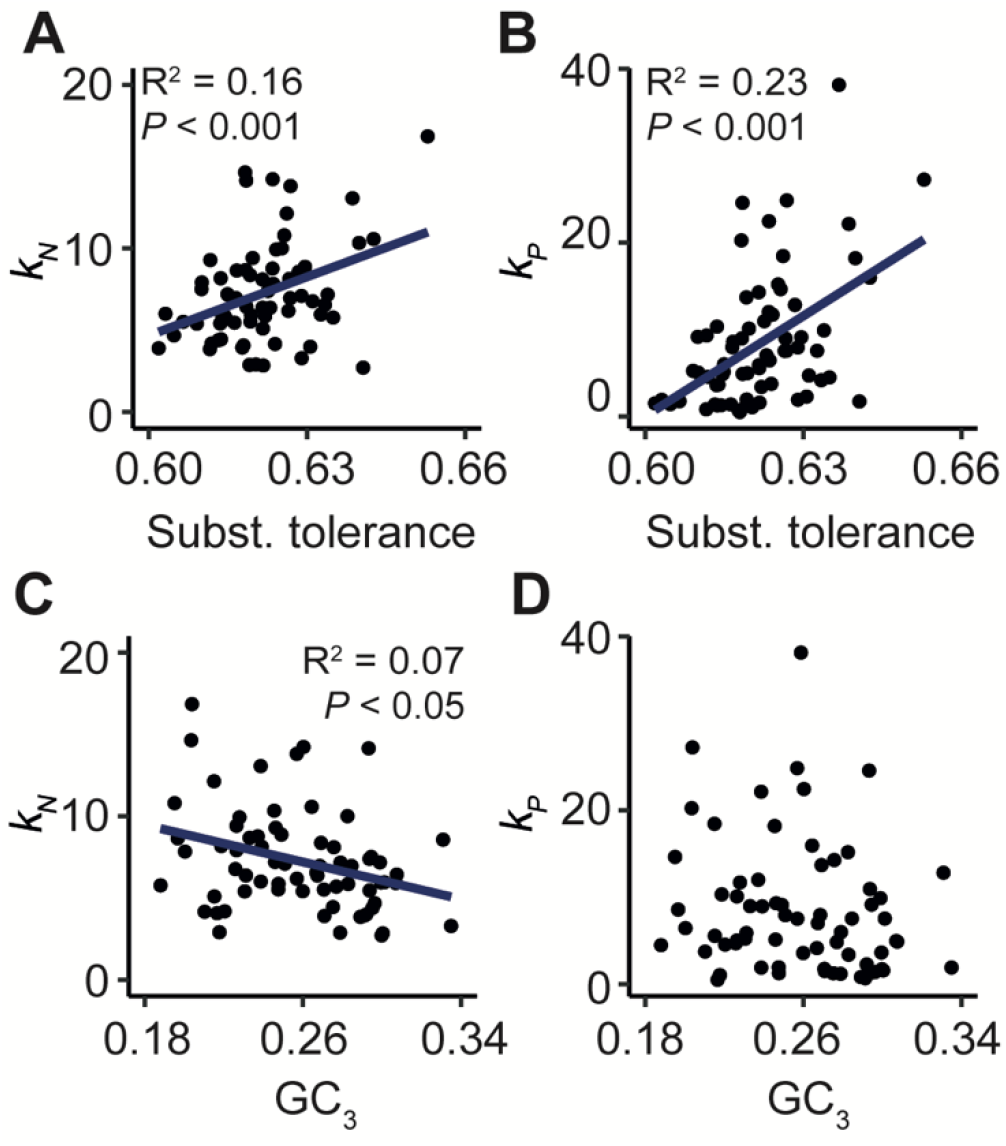
The effect of gene composition on the rate of molecular evolution for plastid encoded genes. The nucleotide composition of a gene was determined by analysing the GC content at the 3^rd^ codon position (GC_3_). Meanwhile, amino acid composition of the encoded protein was summarised in a substitution tolerance score based on the BLOSUM62 matrix, where a lower value corresponds to proteins with a composition less tolerant to substitution. **A-B**) Scatter plots showing protein substitution tolerance scores versus the rate of nucleotide evolution and the rate of protein evolution, respectively. **C-D**) Scatter plots showing GC content at the 3^rd^ codon position (GC_3_) versus the rate of nucleotide and protein evolution, respectively. Where appropriate, linear models are shown in blue with their associated R^2^ and *P*-values.

### The intrinsic nucleotide composition has only weakly influenced the rate of molecular evolution

Given that the composition of the encoded amino acid sequence has contributed significantly to the rate at which the underlying molecular sequence of a gene has evolved, it was next determined whether similar composition-based constraints were manifest at the level of the nucleotide sequence. There is some precedent for hypothesising that this might occur, as genes encoded in the inverted repeat regions have higher guanine-cytosine (GC) contents and have evolved more slowly than genes located in the single-copy regions (Li, et al. 2016). To prevent re-detecting differences in amino acid composition at the nucleotide level, only the GC content of the third codon position (GC_3_) was investigated, excluding amino acids that have no degeneracy at this position. Comparison of the GC_3_ content of a gene to its rate of molecular evolution revealed that the GC_3_ could only explain ∼7% of the variance in the rate of nucleotide evolution (R^2^ = 0.07, *P* < 0.05), where genes with higher GC_3_ contents have evolved more slowly (Figure 3C). Meanwhile, no significant relationship was identified between GC_3_ and the rate of protein evolution (Figure 3D). Thus, while genes that have higher GC_3_ contents had slower evolving nucleotide sequences, this interaction is weak and GC_3_ has not had an impact on the rate of protein evolution.

### The location of a gene in the plastid genome has influenced its rate of molecular evolution

As noted in the introduction, the rate and type of mutation along the plastid genome is non-uniform. Therefore, it is likely that the position of a gene in the plastid genome could influence its rate of molecular evolution. Given that all the genes analysed here are in the single-copy regions of the plastid genome, gene position was assessed by calculating the average distance from each gene’s midpoint to the closest border of an inverted repeat region in all 773 species. This distance was then compared with the rate of molecular evolution of each gene. This revealed that the further away a gene is located from the inverted repeat, the lower its rate of nucleotide evolution (Figure 4A). In total, distance to the inverted repeat border could explain ∼20% of the variance the rate of nucleotide evolution between genes (R^2^ = 0.20, *P* < 0.001). A similar effect was observed for the rate of protein evolution (R^2^ = 0.11, *P* < 0.01) (Figure 4B). This distance effect was also observed if the rate of molecular evolution of individual genes was aggregated at the level of operons (Supplementary Figure 3). Thus, the rate at which a plastid gene has evolved during the radiation of the angiosperms is dependent on how distant that gene is positioned from the inverted repeat in the plastid genome.

**Figure 4.**
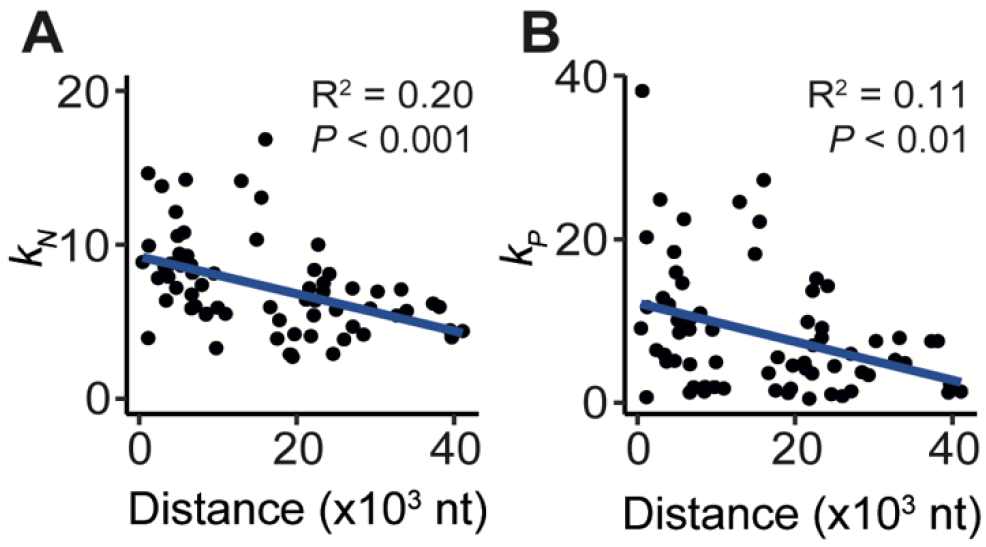
The relationship between the average distance of a gene to the closest inverted repeat border and the rate of molecular evolution of 69 single-copy plastid encoded genes. Scatter plots showing the average distance of each gene’s midpoint to the closest inverted repeat border measured in nucleotides (nt) versus the rate of nucleotide evolution (**A**) and the rate of protein evolution (**B**). Linear models are shown in blue with their associated R^2^ and *P*-values.

### mRNA abundance is a determinant of plastid gene evolutionary rate

High gene expression has been previously linked to low rates of molecular evolution in multiple systems (Pál, et al. 2001; Subramanian and Kumar 2004; Wright, et al. 2004; Seward and Kelly 2018). To investigate whether this phenomenon was also true of genes encoded in the plastid genome, the rate of molecular evolution of a plastid gene was compared to both the mRNA and protein abundance of that gene. In agreement with previous analysis in other systems, genes with high mRNA abundance had low levels of molecular evolution and *vice versa* (Figure 5). Specifically, variance in mRNA abundance could explain 32% of the variation in the rate of nucleotide evolution (R^2^ = 0.32, *P* < 0.001) (Figure 5A) and 27% of variation in the rate protein evolution (R^2^ = 0.27, *P* < 0.001) (Figure 5B), respectively. In contrast, there was no significant relationship between protein abundance and rate of molecular evolution (Figure 5C and 5D). This suggests that the constraint on the evolutionary rate is mediated via an interaction with transcription rather than through the requirement for protein production. Furthermore, previously detected interactions between mRNA abundance and gene evolutionary rate have been manifest in part through selection acting on transcript biosynthesis cost (the number and type of atoms contained within the transcript and the number of high-energy phosphate bonds required biosynthesis) and transcript translation efficiency (a function of the number of tRNAs that can translate that codon encoded in the plastid genome) (Seward and Kelly 2016, 2018). However, there was no evidence for an interaction between these two factors and the rate of molecular evolution of genes encoded in the plastid genome (Supplementary Figures 4 and 5). Thus, the extent to which a transcript accumulates in the chloroplast has constrained the evolutionary rate of its corresponding gene, and this is independent of selection acting to preserve the biosynthetic cost or translational efficiency of the coding sequence or the abundance of the encoded protein.

**Figure 5.**
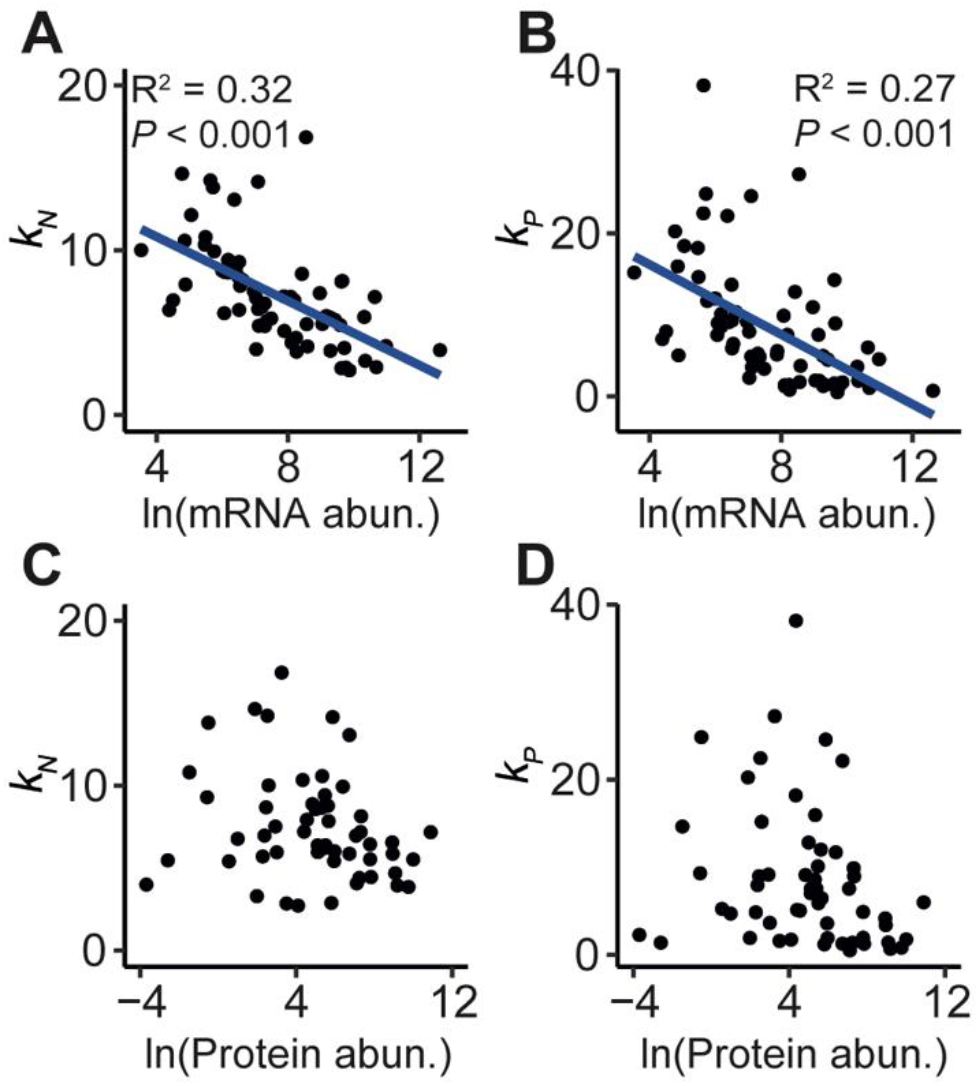
Relationship between the rate of molecular evolution and gene expression for plastid encoded protein-coding genes. **A-B**) Scatterplots of mRNA abundance (data from (Forsythe, et al. 2022)) versus the rate of nucleotide evolution and protein evolution, respectively. Linear regressions are shown as blue lines with their associated R^2^ and *P*-values. **C-D**) Scatterplots of protein abundance (data from (Wang, et al. 2012)) versus nucleotide evolution and protein evolution, respectively.

### Covariate models can explain ≥70% of the variance in the rate of molecular evolution of plastid encoded genes

Given that several factors were identified that could each explain a significant component of variation in the rate of molecular evolution of plastid genes in isolation, we next sought to identify the cumulative variance attributable to all factors while accounting for interdependencies (Figure 8A) which may deflate the explained variance observed when single variables are considered. To do this, regression tests were conducted for all possible subsets of the set of independent variables described above. This revealed that *d*_N_/*d*_S_, gene distance to the inverted repeat border, protein substitution tolerance and mRNA abundance together could explain 70% of the variance in the rate of nucleotide evolution (Figure 6B, Supplementary Figure 6). Relative importance metrics showed *d*_N_/*d*_S_ is the most important covariate (30.7% variation explained), followed by mRNA abundance (16.8% variation explained), gene distance to the inverted repeat (14.3% variation explained) and then protein substitution tolerance (7.8% variation explained), respectively (Figure 6B). Thus, in total 39% of variation in the rate of nucleotide evolution between genes is attributable to compositional factors and production requirements. A similar result was obtained for the proportion of variance explained in the rate of protein evolution (Figure 6C, 78% variance explained, Supplementary Figure 7). Specifically, *d*_N_/*d*_S_ was the most important covariate (46.5% variation explained), followed by mRNA abundance (12.6% variation explained), protein substitution tolerance (11.9% variation explained), and then gene distance to the inverted repeat (7.2% variation explained), respectively (Figure 6B). Thus, in total 32% of variation in the rate of protein evolution between genes is attributable to compositional factors and production requirements. In all cases, the weak association between GC_3_ content and the rate of molecular evolution (Figure 6) was rendered insignificant when other factors above were accounted for.

**Figure 6.**
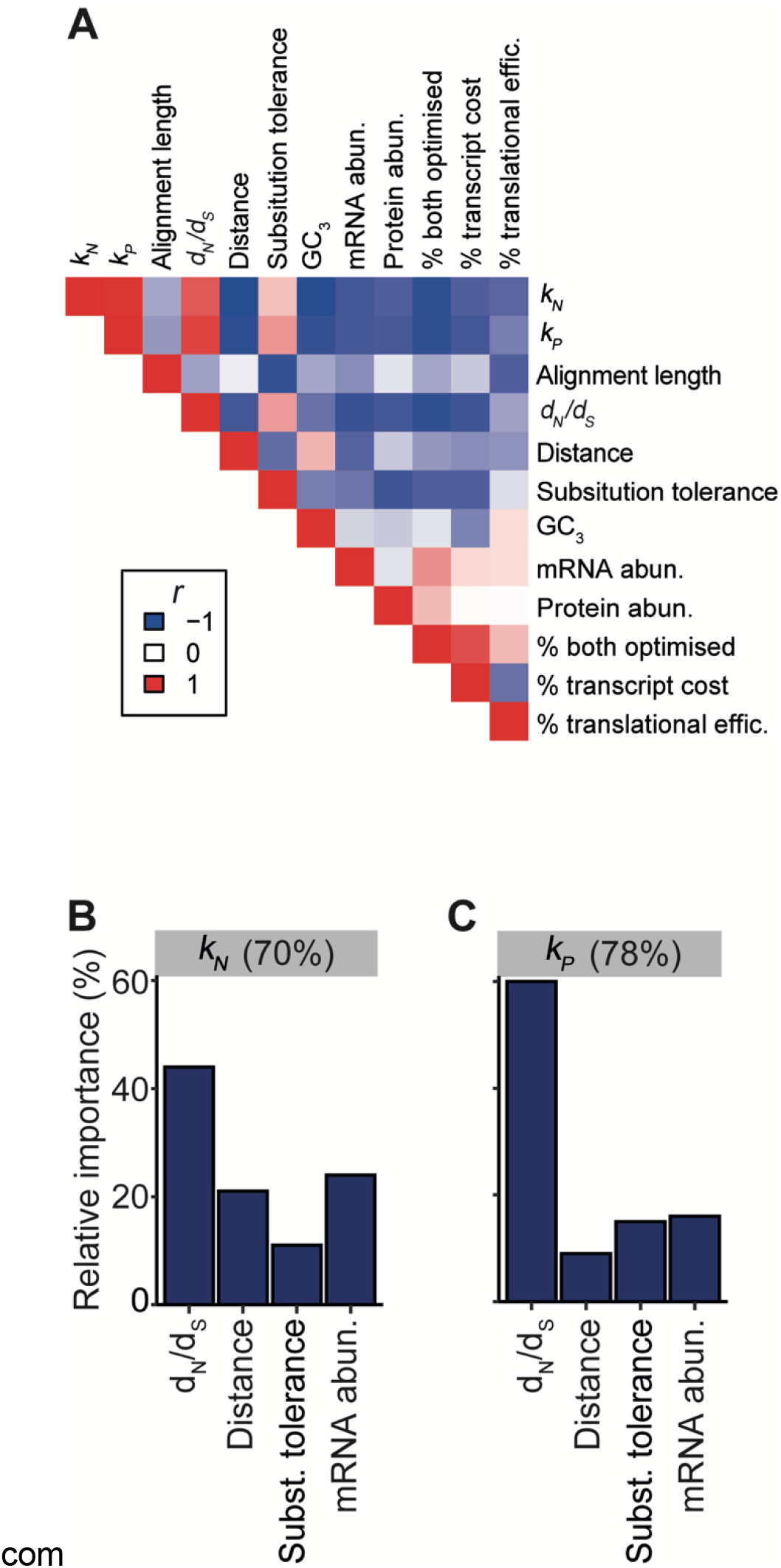
Relative importance of covariates in the best linear models identified to explain the rates of molecular evolution. **A)** A correlation heatmap of all continuous variables used in this study. Red, Pearson correlation coefficient (*r*) = 1; white, *r* = 0; blue, *r* = -1. **B-C)** The relative importance (given as a percentage) of *d*_N_/*d*S, gene distance to the inverted repeat, protein substitution tolerance and mRNA abundance in explaining 70% and 78% of the variance in the rate of nucleotide and protein evolution, respectively.

To ensure model space was effectively explored, random forests were also run on the complete set of factors to determine if there were alternative models that could explain the variance in the rate of molecular evolution (Supplemental File 1). None of the random forest models outperformed the linear regression analyses shown above, despite including more parameters (Supplemental File 1). Thus, only four variables are required to explain ≥70% of the variance in nucleotide and protein evolution in the plastid genome. Moreover, compositional factors and production requirements intrinsic to the genes themselves explain ≥32% of the variation in the rate of molecular evolution and have constrained the capacity for adaptive evolution in the chloroplast genome.

## Discussion

The endosymbiosis of the bacterial progenitor of the chloroplast is a landmark event in the evolution of eukaryotes. It first enabled oxygenic photosynthesis in eukaryotes and helped direct the Earth on a continued trend of decreasing atmospheric CO_2_ and increasing O_2_ over the subsequent ∼1.5 billion years (Holland 2002; Canfield 2005). Following this endosymbiosis there was a dramatic reduction in the gene content of the plastid genome, such that it now harbours fewer than 5% of the genes found in its free-living bacterial relatives (Palmer 1985; Timmis, et al. 2004). These remaining genes have been trapped in the same genome and inherited in the absence of sexual recombination throughout their evolution (Birky 1995; Mogensen 1996). Here, we investigate the factors that have contributed to the variation in evolutionary rate of plastid encoded genes over the last 125 million years during the radiation of the angiosperms. We show that variation in the strength of purifying selection between genes has severely constrained the rate of molecular evolution. We further show that the rate of molecular evolution has also been governed by factors that are extrinsic to the function of the gene. Specifically, the rate of molecular evolution of plastid genes has been influenced by the position of the gene in the plastid chromosome, the substitution tolerance of the encoded protein and the level of gene expression. Combined, the production requirement and compositional factors explain 39% and 32% of the variance in the rate of nucleotide and protein evolution of plastid encoded genes, respectively. Thus, the capacity for molecular evolution of a plastid encoded gene has been determined in part by the location, composition, and expression level of that gene.

It has previously been shown that plastid genes are under strong purifying selection (Drouin, et al. 2008; Smith 2015). Purifying selection is a measure of the functional constraint on a protein sequence, whereby greater functional constraint results in the increased likelihood of an amino acid change having a deleterious effect and subsequently being eliminated from the population. *A priori* one might expect variation in purifying selection between genes to explain most of the variation in rate of protein evolution between genes and have a knock-on effect on modulating the rate of nucleotide evolution. However, the phylogenetically resolved analysis presented here revealed that the strength of purifying selection to which a gene is subject explained 47% and 31% of the rate of protein and nucleotide evolution, respectively. Thus, although differences in the strength of purifying selection is a major component of variation in the rate of molecular evolution, it does not account for all rate variation between genes.

The finding that genes which are positioned further from the inverted repeat border, and hence the likely origin of replication, are evolving more slowly than genes that are closer has two of important implications. First, this finding is compatible with models of chloroplast DNA replication where replication initiates at origins located in the inverted repeat, and where DNA closer to the inverted repeat border spends more time in a single-stranded state than DNA that is more distant to the origin. This is because single-stranded DNA is more susceptible to base deamination which can result in mutation and elevated rates of evolution (Lindahl 1993; Krishnan, et al. 2004). Thus, if DNA closer to the origin of replication spends more time single-stranded during replication, this could lead to a positional effect where genes closer to the inverted repeat have elevated rates of molecular evolution. One model of plastid genome replication that satisfies both these criteria is the bi-directional dual displacement loop mechanism (Kolodner and Tewari 1975). However, there is increasing evidence that plastid genome replication proceeds via a recombination dependent mechanism (Oldenburg and Bendich 2004; Oldenburg and Bendich 2016), and while these mechanisms have also been proposed to initiate in the inverted repeat, it is difficult to conceptualise how DNA would spend differential time single-stranded in these models. Therefore, our findings are consistent with the bi-directional dual displacement loop mechanism and lend further support to analyses of intergenic regions and the third codon position that indicate the wide-spread operation of this form of plastid DNA replication in angiosperms (Krishnan and Rao 2009). Second, the fact that origin-proximal genes are evolving faster than origin-distal genes means that all locations in the plastid genome are not equal in terms of their capacity to generate genetic diversity. Thus, the position of a gene and the structure of the plastid genome has constrained the evolvability of adaptive changes in the chloroplast.

The 20 amino acids that are used to build proteins vary in the physical and chemical properties of their side chains: some are hydrophobic, some are charged, some are large, and some are small. Consequently, a mutation that results in the substitution of one amino acid for another with similar physicochemical properties is more likely to have a smaller impact on the structure and function of a protein than substitution with an amino acid with substantially different properties (Sneath 1966; Epstein 1967; Henikoff and Henikoff 1992; Bohórquez, et al. 2017). As there are different numbers of similar amino acids in each physicochemical category, and each amino acid can occupy different physicochemical categories, each amino acid has a different tolerance to substitution. We estimated the substitution tolerance of each protein sequence encoded in the plastid genome and found that the aggregate substitution tolerance of a protein sequence is a major component of variation in its evolutionary rate. This means that the evolvability of novel adaptive phenotypes arising from variation in plastid encoded genes is limited by the identity of the encoded amino acids.

In agreement with previous analyses in other systems, we observed that plastid genes with high expression levels had lower rates of molecular evolution (Pál, et al. 2001; Subramanian and Kumar 2004; Wright, et al. 2004; Seward and Kelly 2018). However, in contrast to previous studies, this phenomenon was not attributable to selection acting to maintain a low transcript biosynthesis cost or a high translational efficiency, nor was it attributable to the requirement for high protein abundance. The lack of association with protein abundance could be due to increased noise in the data as it is an integrated dataset, or alternatively, protein abundance is not a good proxy for protein production as protein turnover is not accounted for. Nevertheless, the results presented here raise the question as to what phenomenon could give rise to a strong interaction between mRNA abundance with the rate of molecular evolution that is not attributable to transcript biosynthetic cost, translational efficiency or protein abundance. One mechanism that could potentially explain this dependency is the presence of transcription mediated DNA repair, whereby increased transcriptional activity directly enhances DNA repair resulting in reduced rates of mutation and molecular evolution. This would give rise to a phenomenon whereby the more highly transcribed a gene, the more likely DNA damage is repaired in that gene and the lower its mutation rate. Interestingly, this potential explanation is supported by the presence of a nuclear-encoded, chloroplast-targeted ortholog of mutation frequency decline protein (Mfd), alternatively named transcription-repair coupling factor (TRCF) (Hays 2002). In bacteria, Mfd facilitates DNA repair at stalled RNA polymerase complexes by recruiting the nucleotide excision repair machinery (Selby and Sancar 1993). Additionally, it is proposed to play a role in promoting homologous recombination mediated repair (Ayora, et al. 1996). While plants lack orthologs of other nucleotide excision repair genes (Hays 2002), they contain orthologs of the recombination mediated repair factors (Maréchal and Brisson 2010) and recombination-based repair is a primary DNA repair mechanism in the plastid (Cerutti, et al. 1995; Khakhlova and Bock 2006; Maréchal and Brisson 2010). Thus, it is possible that Mfd in plants functions to promote recombination-based repair in response to DNA damage-stalled RNA polymerases. This effect would likely arise from Mfd’s ability to form R-loops by distorting the DNA around the stalled polymerase (Portman, et al. 2021). This, in turn, is known to lead to recruitment of plastid homologous recombination machinery to facilitate DNA repair (Wang, et al. 2021). Thus, although it is impossible to prove the existence of transcription-coupled DNA repair from the data presented here, it is consistent with the phenomenon. Furthermore, it will be interesting to determine whether loss of function of Mfd in plants leads to increased mutation in a manner that is consistent with a reduction in transcription mediated DNA repair, and whether it plays a role in the transcription dependent constraint on the rate of molecular evolution of plastid genes.

Our finding that the expression, location, and amino acid composition of a gene impose significant constraints on rate of molecular evolution of plastid encoded genes is important. It means that factors other than the function of the encoded protein have been important in constraining the evolvability of plastid genes. In this context, it is noteworthy that the constraints identified here may help provide a mechanistic explanation for the strong phylogenetic constraints that have limited the adaptation of rubisco enzyme kinetics (Bouvier, et al. 2021) and why rubisco is one of the slowest evolving enzymes on earth (Bouvier, et al. 2022). Specifically, evolutionary constraints intrinsic to the location, mRNA abundance and amino acid composition of rubisco (in addition to functional constrains on the protein) may help to explain why it has been so slow to adapt to changing atmospheric conditions, and consequently why it is poorly suited to the O_2_-rich, CO_2_-poor atmosphere of the present day. Furthermore, it is tempting to speculate that endosymbiotic gene transfer to the nuclear genome allowed other plastid encoded genes to escape some of these evolutionary constraints, facilitated more rapid adaptation of chloroplast proteins, and provided a further advantage for migration of organellar genes to the nuclear genome.

In summary, the molecular adaptation of the land plant chloroplast has been constrained by the location, composition and expression of its genes. The presence of these evolutionary constraints limits the evolvability of adaptive phenotypes arising from chloroplast encoded genes, and thus motivates strategies to improve photosynthesis in synthetic biology contexts by circumventing these evolutionary barriers to optimisation.

## Methods

### Dataset curation

The complete set of fully sequenced plastid genomes was downloaded from the National Center of Biotechnology and Information (NCBI) (https://www.ncbi.nlm.nih.gov/) in July 2021. Those genomes that did not contain a full set of canonical plastid encoded photosystem I, photosystem II, ATPase, NAD(P)H dehydrogenase-like, cytochrome b_6_f complex, ribosomal and RNA polymerase genes were discarded. All other plastid encoded genes were not included as they were missing from a large number of angiosperm plastid genomes (Supplemental Table S2). The full set of genes included in the analysis and their accession numbers are provided in Supplemental File 2. The protein sequences encoded by these genes were aligned using MAFFT L-INS-I (Katoh and Standley 2013), and visually checked in AliView (Larsson 2014).

### Phylogenetic tree inference

The protein multiple sequence alignments were used to guide codon alignment using Pal2Nal (Suyama, et al. 2006) so that there were no inconsistencies between the protein and nucleotide alignments. Nucleotide and protein multiple sequence alignments were trimmed to remove columns containing >90 % gaps using trimAl (Capella-Gutiérrez, et al. 2009). All 69 trimmed nucleotide multiple sequence alignments were concatenated together and used to identify the best-fitting model of sequence evolution using the inbuilt ModelFinder function in IQ-TREE (Kalyaanamoorthy, et al. 2017). In total, 243 models of sequence evolution were tested and the best-fitting model according to the Bayesian information criterion (BIC) was GTR+F+R7. The best-fitting model of protein sequence evolution was identified in a similar manner, with 447 models tested and JTT+F+R7 selected according to BIC. A maximum-likelihood phylogenetic tree was then inferred from the concatenated nucleotide alignment and the best-fit model GTR+F+R7 using IQ-TREE’s ultrafast bootstrapping method with 1000 replicates (Hoang, et al. 2018). The resulting tree was then rooted manually in Dendroscope using the Nymphaeales species as the outgroup as they were the most basal-branching clade in the tree (Huson and Scornavacca 2012).

### Evolutionary rates and selection pressure

To estimate the rate of evolution for each individual gene, gene trees were inferred from the trimmed nucleotide and protein multiple sequence alignments. Here, the topology of the tree was constrained to that identified form the concatenated alignment above. This was done as this is the topology that is most likely to be the true topology for all genes given the uniparental inheritance of the plastid in the absence of sexual recombination – i.e. all gene trees should have the same topology. The model of sequence evolution was also constrained to that obtained using the concatenated alignments as this is most likely to be correct, and holding this constant enables comparison of inferred rates of molecular evolution between genes. Gene trees were inferred with the above constraints using IQ-TREE (Nguyen, et al. 2015). To perform an error check and ensure that the topology of the individual gene trees was identical to the species tree, the Robinson-Foulds distance was calculated (Sukumaran and Holder 2010). For each gene tree, the total tree length, which was the sum of all the branch lengths, was extracted from the IQ-TREE output and used as a measure of the rate of sequence evolution (*k*). The ratio of the rate of non-synonymous to synonymous mutation rate (*d*_N_/*d*_S_) was calculated using IQ-TREE from gene trees constrained to the topology of the tree generated from the concatenated nucleotide sequence using the codon model of sequence evolution MGK+F3×4. There were too few indels to facilitate an analysis of the rate of index evolution (Supplemental File 3).

### Protein tolerance to substitution

Protein tolerance to substitution scores were calculated using the BLOSUM62 matrix (Henikoff and Henikoff 1992). BLOSUM62 was chosen as it is the most widely used substitution matrix the biological sciences, being used in BLAST (and related sequence search methods) and nearly all protein multiple sequence alignment methods. Here, the odds ratios for all substitutions were back calculated from the log-odds scores provided. Each residue was then given a substitution tolerance by calculating the geometric mean of the odds ratios of its substitution to the remaining 19 common residues. Then, for each of the 69 proteins analysed, the average residue substitution tolerance was calculated by taking the arithmetic mean of the concatenated protein sequence (composed from all 773 species sequences for that protein).

### Gene position and GC_3_ content

Although the plastid genome is a highly conserved structure in the angiosperm lineage, its size, and thus gene position, varies between species. Since all of the genes included in this analysis are in the single-copy region, the average genetic distance of a gene’s midpoint to the closest inverted repeat border was used as a measure of gene position relative to the putative origin of replication (the inverted repeat). To do this, the positions of the inverted repeat regions were identified for each species by searching for large-duplicated regions using BLASTN (Altschul, et al. 1990; Camacho, et al. 2009). Then, the distance between the gene’s midpoint (obtained from the GenBank file for each species’ plastid genome) and the closest inverted repeat border was calculated. An average distance was then determined and this average for each gene was used in the analysis of gene position. Positional effects were also analysed at the level of operons. To do this, operon maps were obtained from (Shahar, et al. 2019) for six diverse species in our dataset. The operon’s distance to the closest inverted repeat border was determined by averaging the distances of its encoded genes.

The average GC content of the 3^rd^ codon position (GC_3_) for all 773 species was calculated for each gene. The GC_3_ excluded methionine and tryptophan codons as these amino acids are encoded by a single codon, and thus have no flexibility in base identity at the 3^rd^ position.

### mRNA abundance and protein abundance

mRNA abundance data were obtained from (Forsythe, et al. 2022) and values were available for all 69 genes analysed in this study. The whole organism integrated (3702-WHOLE_ORGANISM-integrated.txt) protein abundance estimates for *Arabidopsis thaliana* were obtained from the Protein Abundance Database (PaxDb) (Wang, et al. 2012). Abundance data was available for 56/69 genes included in this analysis.

### Transcript biosynthetic cost and translational efficiency

The strength of selection acting to reduce transcript cost, increase translational efficiency and the trade-off between these two forces was inferred for each species using CodonMuSe (Seward and Kelly 2018). The tRNA copy numbers required by CodonMuSe were inferred for each species by inputting the species plastid genome to tRNAscan-SE 2.0 (Chan, et al. 2021).

### Statistical analysis and modelling

All statistical analysis and linear regressions were calculated in R. Identification of the best subset of variables for multiple linear regressions was determined using the olsrr package. Relative importance metrics were generated using the reliampo package (Groemping 2006). Random forests were using the randomForest package (Svetnik, et al. 2003).

## Acknowledgements

This work was funded by the Royal Society and the European Union’s Horizon 2020 research and innovation program under grant agreement number 637765. ER is funded by the Biotechnology and Biological Sciences Research Council (BBSRC) [grant numbers BB/M011224/1 and BB/P003117/1]. This research was funded in whole, or in part, by the BBSRC. For the purpose of open access, the author has applied a CC BY public copyright license to any Author Accepted Manuscript version arising from this submission.

## Data Availability

All data used in this study is provided in the supplemental material, and all accession numbers are provided.

## Author Contributions

ER and SK conceived the study. ER conducted the analysis. ER and SK wrote the manuscript.

