## Supplemental File 1 for "The evolutionary constraints on angiosperm chloroplast adaptation"

***Supplementary File 1***

Random forest regressions

Random forest regressions were used to see how much variance in the rate of nucleotide evolution and protein evolution could be explained by the variables explored in this study. The input for the random forest regressions included $d_N/d_S$, gene distance to the inverted repeat, mRNA abundance, protein abundance, % optimised for transcript biosynthetic cost, % optimised for translational efficiency, % optimised for the trade-off between transcript cost and translational efficiency, protein substitution tolerance (see methods) and gene $GC_3$ content.

*Rate of nucleotide evolution*

The percentage of variance in the rate of nucleotide evolution explained by the random forest model using all input variables was 56% (mean squared of residuals = 5.26). This is less than the variance explained by the best multiple linear regression model (70%, see results) which contained only 4 covariates, $d_N/d_S$, gene distance to the inverted repeat border, protein substitution tolerance and mRNA abundance. However, in agreement with our linear models, the random forest regression identified these four co-variates as four of the five most important variables (Table 1, Figure 1).

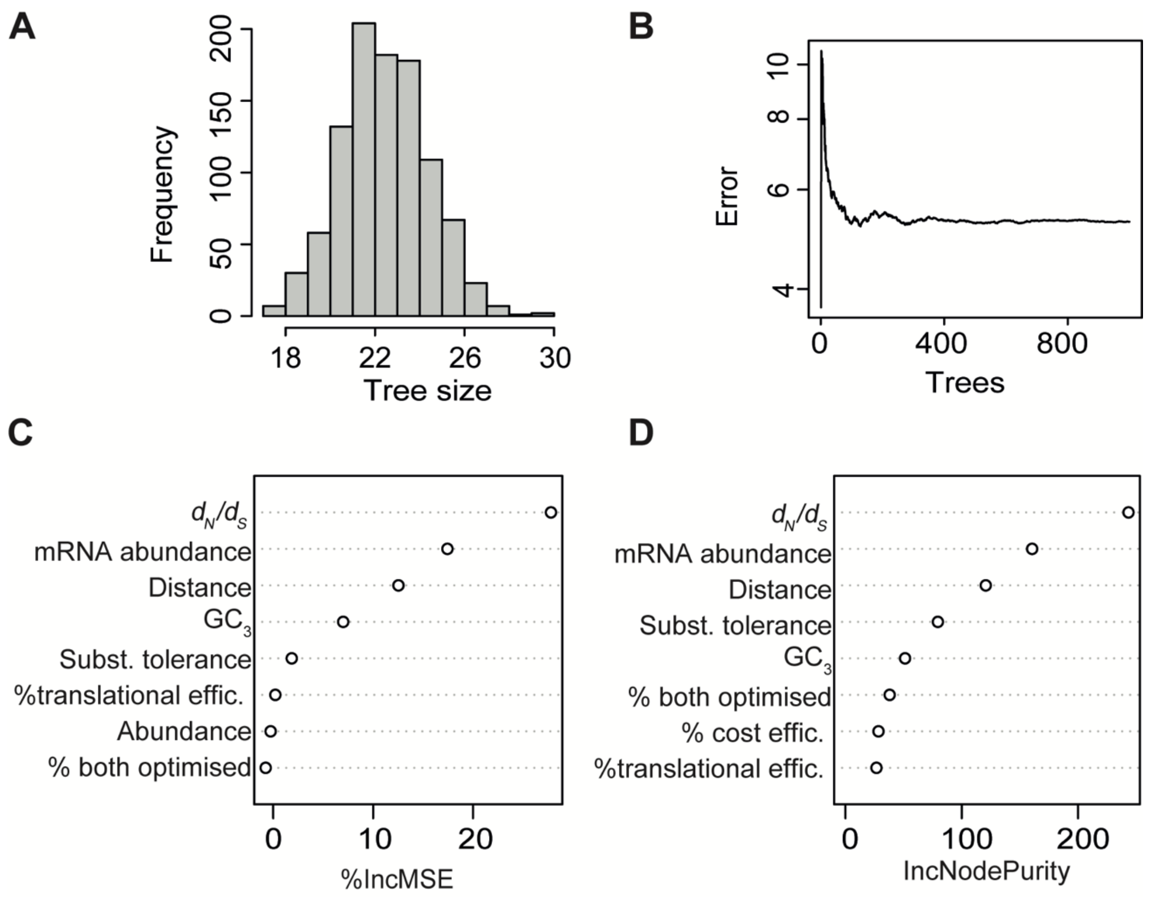


**Figure 1.** Random forest regression for the rate of nucleotide evolution. **A)** Histogram of tree size for the 1000 trees generated. **B)** Error rate versus the number of trees built. **C)** The percentage increase in mean squared error (%IncMSE) for all variables included in the regression ordered in decreasing value from top to bottom. **D)** The increase in node purity (IncNodePurity) in decreasing value from top to bottom.

**Table 1.** Variable importance metrics for random forest regression of the rate of nucleotide evolution.

| Variable | Mean minimum depth1 | No. of nodes2 | MSE increase3 | Node purity increase4 | No. of trees5 | No. times a root6 | p-value | adjusted p-value |
|---|---|---|---|---|---|---|---|---|
| $d_N/d_S$ | 1.36 | 3464 | 5.03 | 243 | 995 | 276 | 4.54E-97 | 4.09E-96 |
| mRNA abundance | 1.68 | 3193 | 2.25 | 161 | 979 | 253 | 7.28E-55 | 3.28E-54 |
| Distance | 2.10 | 2912 | 1.36 | 121 | 965 | 178 | 2.07E-23 | 6.21E-23 |
| substitution tolerance | 2.77 | 2479 | 0.16 | 79 | 936 | 127 | 1.94E-01 | 4.36E-01 |
| Abundance | 4.10 | 1615 | -0.01 | 20 | 795 | 9 | 1.00E+00 | 1.00E+00 |
| % Both optimised | 3.46 | 1967 | -0.05 | 38 | 874 | 71 | 1.00 | 1.00E+00 |
| % cost optimised | 3.55 | 2082 | -0.04 | 28 | 884 | 14 | 1.00 | 1.00E+00 |
| % translational optimised | 3.58 | 2013 | 0.01 | 26.68 | 866 | 8 | 1.00 | 1.00E+00 |
| GC3 | 3.021704 | 2220 | 0.50402295 | 51.28938 | 902 | 64 | 1.00E+00 | 1.00E+00 |

1 Mean depth of the node that first splits the data using on that variable.
2 Number of nodes that are split based on that vaiable
3 Increase in mean squared error when that variable is excluded from the decision tree.
4 Increase in node purity from splitting on that variable
5 Number of trees in which that variable is used
6 Number of times that variable is used to split the data at the root of the decision tree

*Rate of protein evolution*

The percentage of variance in the rate of protein evolution explained by the random forest model using all input variables was 67% (mean squared of residuals = 18.9). As for the rate of nucleotide evolution, this was also less than the variance explained by the best multiple linear regression model (78%, see results) which contained 4 covariates, $d_N/d_S$, gene distance to the inverted repeat protein substitution tolerance and mRNA abundance. However, in agreement with our linear models, the random forest regression identified these four co-variates as four of the five most important variables (Table 3, Figure 2).

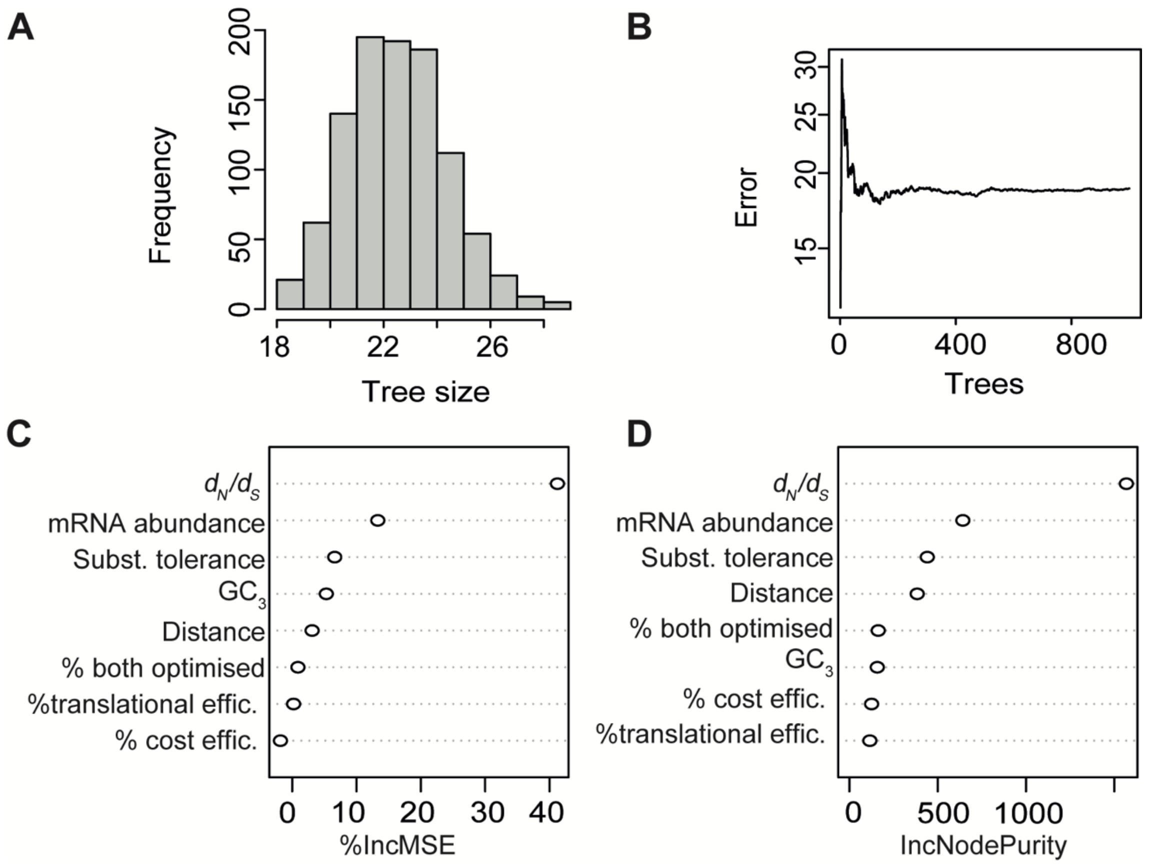


**Figure 2.** Random forest regression for the rate of protein evolution. **A)** Histogram of tree size for the 1000 trees generated. **B)** Error rate versus the number of trees built. **C)** The percentage increase in mean squared error (%IncMSE) for all variables included in the regression ordered in decreasing value from top to bottom. **D)** The percentage node purity (IncNodePurity) in decreasing value from top to bottom.

**Table 2.** Variable importance metrics for random forest regression of the rate of protein evolution.

| Variable | Mean minimum depth[1] | No. of nodes[2] | MSE increase[3] | Node purity increase[4] | No. of trees[5] | No. times a root[6] | p- value | adjusted p-value |
|---|---|---|---|---|---|---|---|---|
| dNdS | 1.20 | 4283.00 | 40.06 | 1568.49 | 996.00 | 342.00 | 0.00 | 0.00 |
| mRNA abundance | 1.87 | 2995.00 | 7.00 | 642.31 | 973.00 | 231.00 | 0.00 | 0.00 |
| Abundance | 3.81 | 1815.00 | -0.52 | 89.92 | 845.00 | 7.00 | 1.00 | 1.00 |
| Distance | 2.56 | 2416.00 | 1.22 | 383.90 | 923.00 | 135.00 | 0.73 | 1.00 |
| % both optimised | 3.54 | 1887.00 | 0.23 | 163.11 | 863.00 | 81.00 | 1.00 | 1.00 |
| % cost optimised | 3.65 | 1935.00 | -0.52 | 125.62 | 875.00 | 18.00 | 1.00 | 1.00 |
| % translational eff. optimised | 3.62 | 2036.00 | 0.04 | 115.69 | 883.00 | 11.00 | 1.00 | 1.00 |
| substitution tolerance | 2.52 | 2450.00 | 2.35 | 441.03 | 947.00 | 148.00 | 0.45 | 1.00 |
| $GC_3$ | 3.21 | 2178.00 | 1.34 | 157.88 | 899.00 | 27.00 | 1.00 | 1.00 |

1 Mean depth of the node that first splits the data using on that variable.

2 Number of nodes that are split based on that vaiable

3 Increase in mean squared error when that variable is excluded from the decision tree.

4 Increase in node purity from splitting on that variable

5 Number of trees in which that variable is used

6 Number of times that variable is used to split the data at the root of the decision tree
