## Supplemental Figures for "The evolutionary constraints on angiosperm chloroplast adaptation"

*Supplementary Figures*

*Supplementary Figure 1*

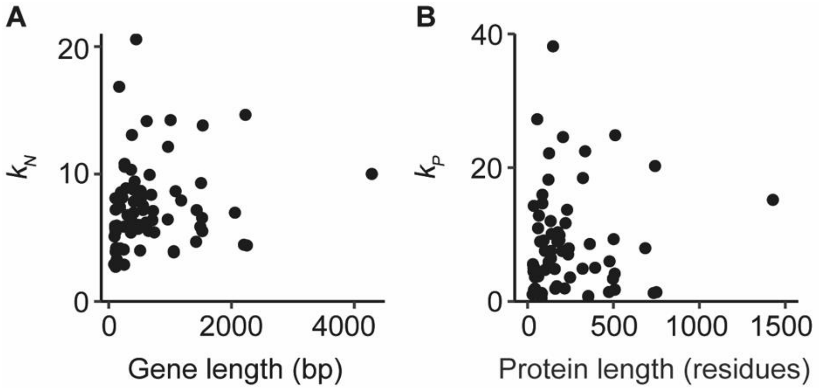


**Supplementary Figure 1.** Relationships between the rates of molecular evolution and sequence length for 69 plastid-encoded genes. **A**) Rate of nucleotide evoution ($k_N$) versus average gene length given in base pairs (bp). **B**) Rate of protein evolution ($k_P$) versus the average protein length given in residues. Rates of molecular evolution have the units of the total substitutions per sequence site per tree ($S_{st}$).

***Supplementary Figure 2***

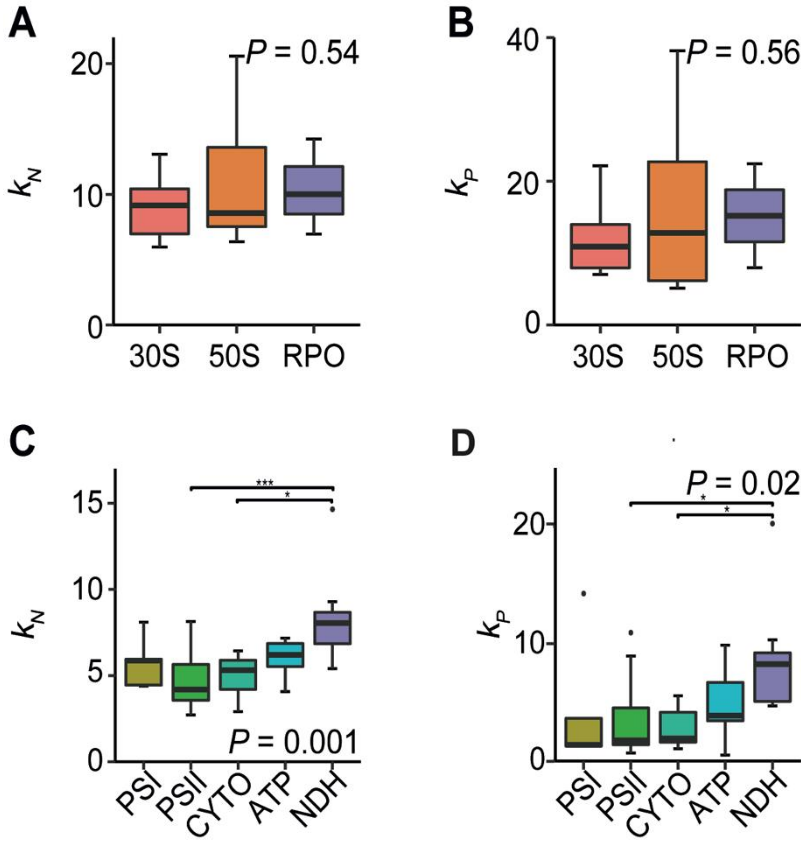


**Supplementary Figure 2.** Exploring the differences in rates of molecular evolution between the multiprotein complexes encoded by the plastid. The difference in the rate of nucleotide evolution and the rate of protein evolution between the major multiprotein complexes of the information processing category (**A** and **B**) and energy production category (**C** and **D**), respectively. Groups include the large ribosomal subunit (50S, n = 7), small ribosomal subunit (30S, n= 10), RNA polymerase (RPO, n = 3), photosystem I (PSI, n = 5), photosystem II (PSII, n = 15), cytochrome $b_6$f complex (CYTO, n = 6), ATP-synthase (ATP, n = 6) and NADPH dehydrogenase-like complex (NDH, n = 10). Statistical significance was assessed using one-way ANOVAs and their associated *P*-values are displayed. Where appropriate, asterisks indicate the *P*-value significance levels of *post-hoc* Tukey tests.

***Supplementary Figure 3***

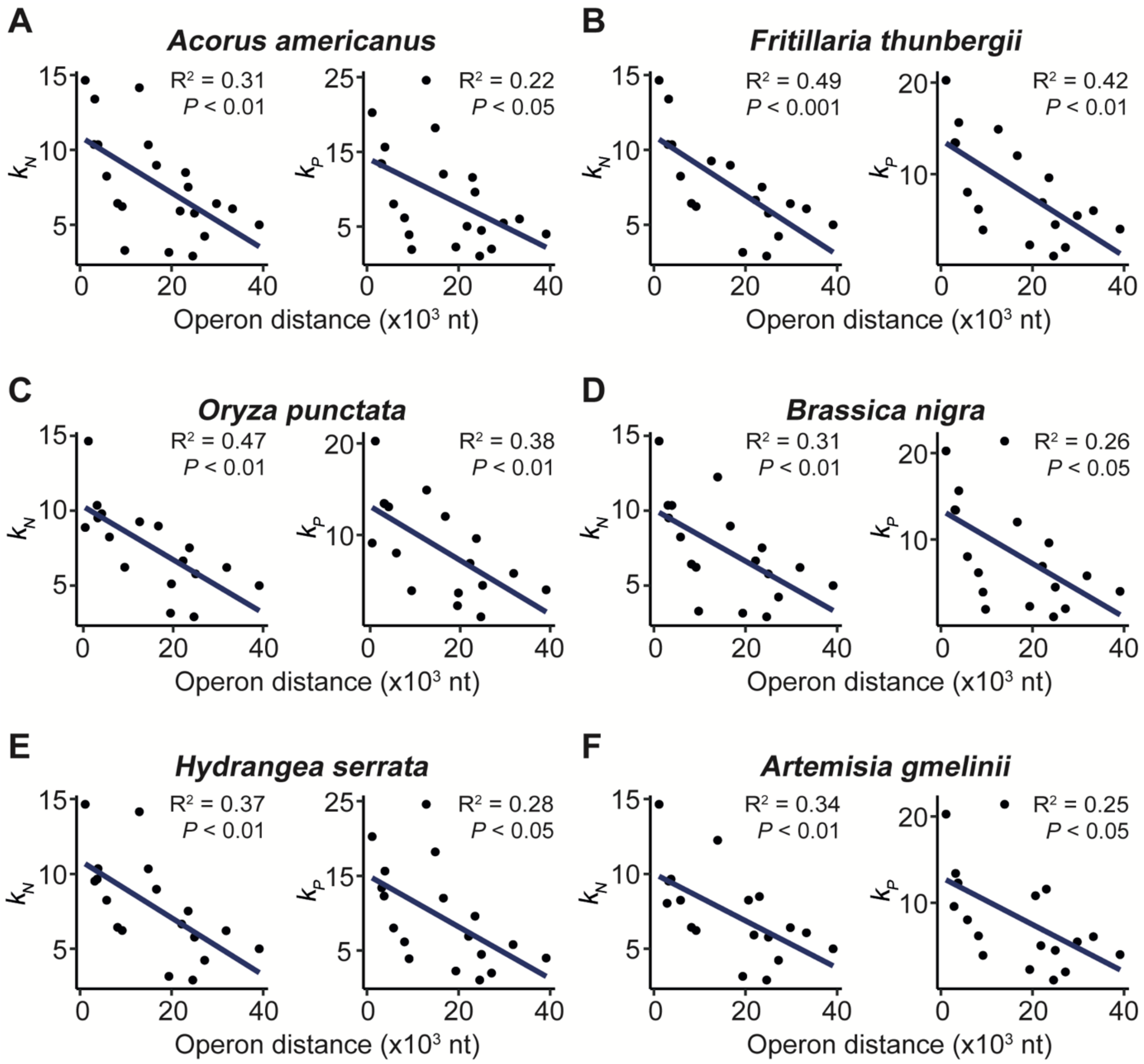


**Supplementary Figure 3.** The relationship between the distance of an operon to the closest inverted repeat border and the rate of molecular evolution of its encoded genes for six species from across the tree analysed in this study. These species included three monocots and three eudicots, all from different orders. The rate of molecular evolution of an operon is calculated from the average rate of molecular evolution of its encoded genes that were included in this analysis. Scatter plots showing the average distance of each operon to the closest inverted repeat border measured in nucleotides (nt) versus the rate of nucleotide evolution (left) and protein evolution (right) for *Acorus americanus* (**A**), *Fritillaria thunbergia* (**B**), *Oryza punctata* (**C**), *Brassica nigra* (**D**), *Hydrangea serrata* (**E**) and *Artemisia gmelinii* (**F**). Linear models are shown in blue with their associated $R^2$ and *P*-values.

***Supplementary Figure 4***

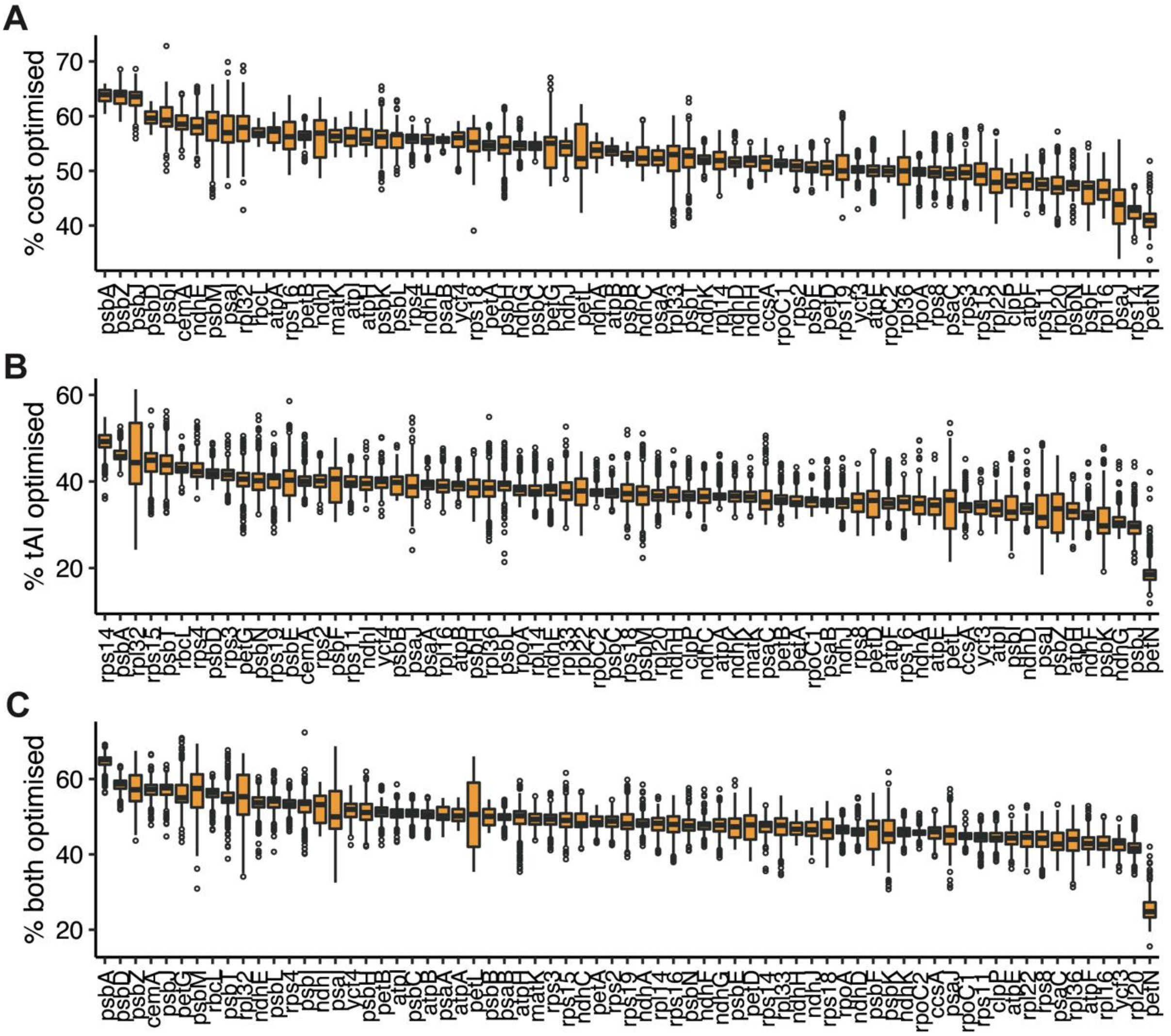


**Supplementary Figure 4.** Extent of gene optimisation for transcript biosynthetic cost and translational efficiency for 69 plastid-encoded genes. For each gene, CodonMuse was used to calculate percent optimisation values for transcript biosynthetic cost (**A**), translational efficiency (**B**) and the trade-off between these two evolutionary forces (**C**) for 773 species which are represented as boxplots. Genes are arranged with decreasing median values from left to right.

***Supplementary Figure 5***

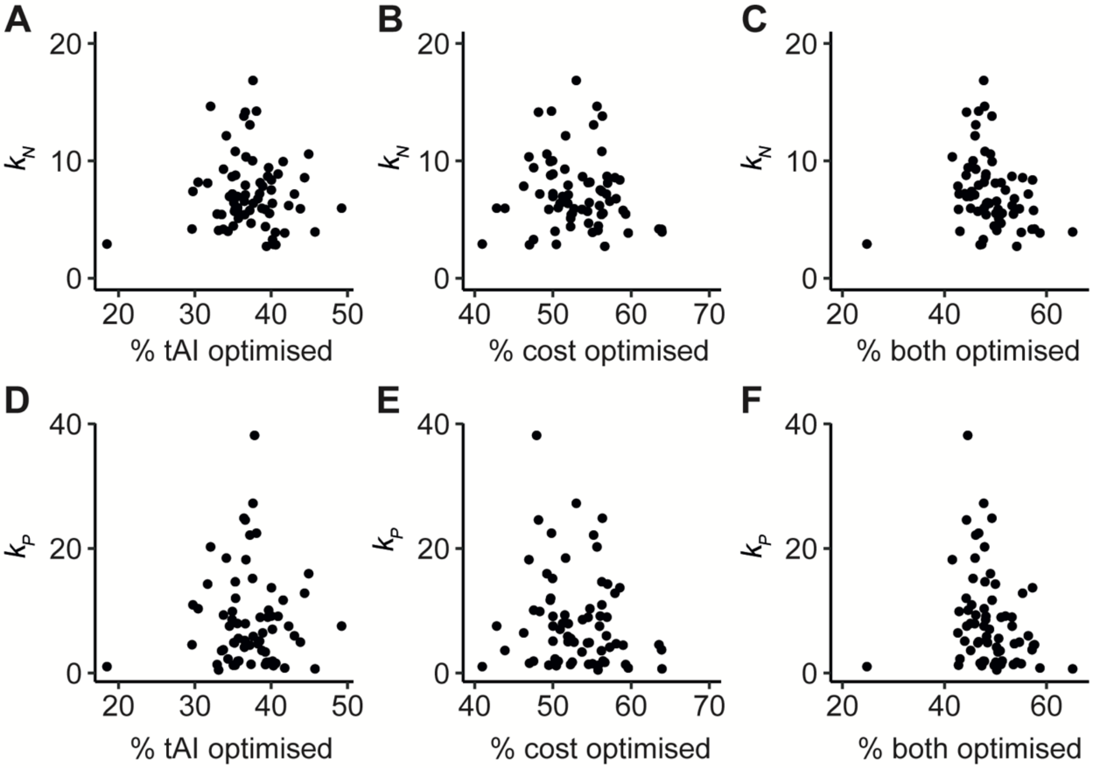


**Supplementary Figure 5.** The relationship between the extent of optimisation of transcript cost and translational efficiency with the rate of molecular evolution. Optimisation values were calculated using CodonMuSe and are given as a percentage, where 100% is the most optimised nucleotide sequence for a given protein sequence. The translational efficiency optimisation score is based on the tRNA adaptation index (tAI) and the cost optimisation scores are based on codon nitrogen cost. Relationships between the median of gene optimisation for translational efficiency, transcript cost and the trade-off with the rate of nucleotide evolution (**A-C**) and protein evolution (**D-F**)

***Supplementary Figure 6***

**A**

| Model Index | Predictors |
|---|---|
| 1 | $d_N/d_S$ |
| 2 | av. IR distance, $d_N/d_S$ |
| 3 | av. IR distance, ln(mRNA abundance), $d_N/d_S$ |
| 4 | av. IR distance, substitution tolerance, ln(mRNA abundance), $d_N/d_S$ |
| 5 | av. IR distance, substitution tolerance, $GC_3$, ln(mRNA abundance), $d_N/d_S$ |
| 6 | av. IR distance, substitution tolerance, $GC_3$, ln(mRNA abundance), $d_N/d_S$, % cost optimised |
| 7 | av. IR distance, substitution tolerance, $GC_3$, ln(mRNA abundance), $d_N/d_S$, % both optimised, % cost optimised |
| 8 | av. IR distance, substitution tolerance, $GC_3$, ln(mRNA abundance), $d_N/d_S$, % both optimised, % cost optimised, % tAI optimised |

**B**

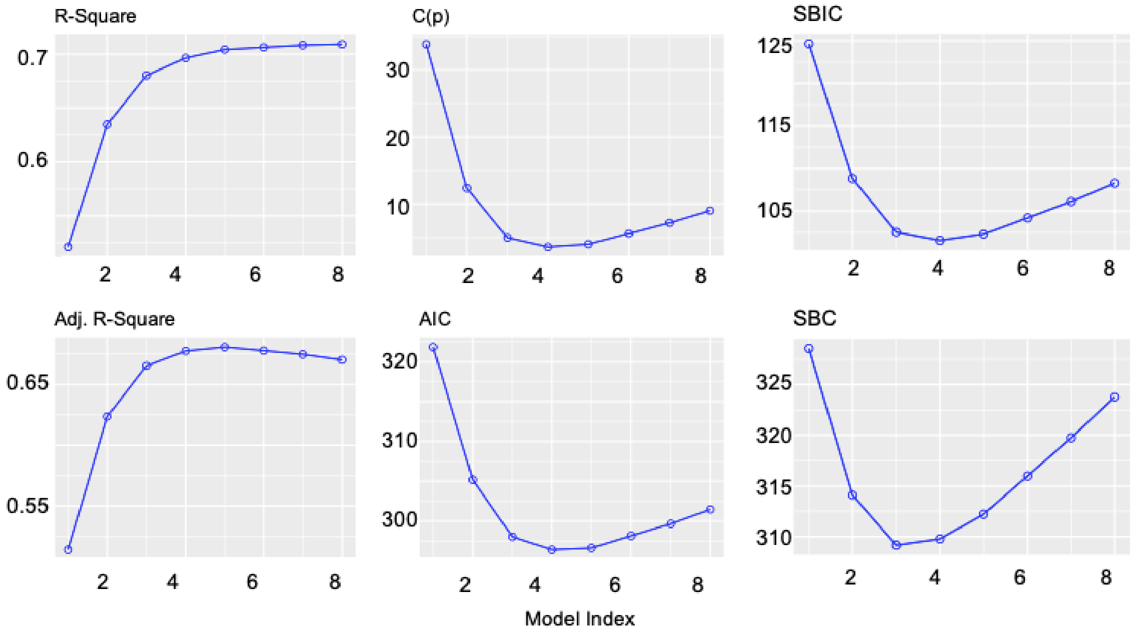


**Supplementary Figure 6.** Variable selection for the best-fitting linear model explaining the maximum variation in the rate of nucleotide sequence evolution. **A**) Table showing the subset of variables used for the different models tested (indexed 1-8). Variables include the non-synonymous to synonymous mutation rate, $d_N/d_S$; the natural logarithm of transcript abundance, ln(mRNA abundance); average gene distance to the inverted repeat, av. IR distance; protein tolerance to substitution, substitution tolerance; mean percent optimisation for translational efficiency, % tAI median; mean percent optimisation for transcript cost, % cost median; mean trade-off for optimisation of both translational efficiency and transcript cost, % both median and GC content of the wobble position, $GC_3$. **B**) Various objective criteria values for the models described in A. R-Square, R-squared of the model; Adj. R-Square, R-squared after adjusting for the number of parameters; C(p), Mallow's Cp; AIC, Akaike information criterion; SBIC, Sawa’s Bayesian information criteria; SBC, Schwarz Bayesian information criteria.

### Supplementary Figure 7

**A**

| Model Index | Predictors |
|---|---|
| 1 | $d_N/d_S$ |
| 2 | av. IR distance, $d_N/d_S$ |
| 3 | av. IR distance, substitution tolerance, $d_N/d_S$ |
| 4 | av. IR distance, substitution tolerance, ln(mRNA abundance), $d_N/d_S$ |
| 5 | av. IR distance, substitution tolerance, $GC_3$, ln(mRNA abundance), $d_N/d_S$ |
| 6 | av. IR distance, substitution tolerance, $GC_3$, ln(mRNA abundance), $d_N/d_S$, % cost optimised |
| 7 | av. IR distance, substitution tolerance, $GC_3$, ln(mRNA abundance), $d_N/d_S$, % both optimised, % cost optimised |
| 8 | av. IR distance, substitution tolerance, $GC_3$, ln(mRNA abundance), $d_N/d_S$, % both optimised, % cost optimised, % tAI optimised |

**B**

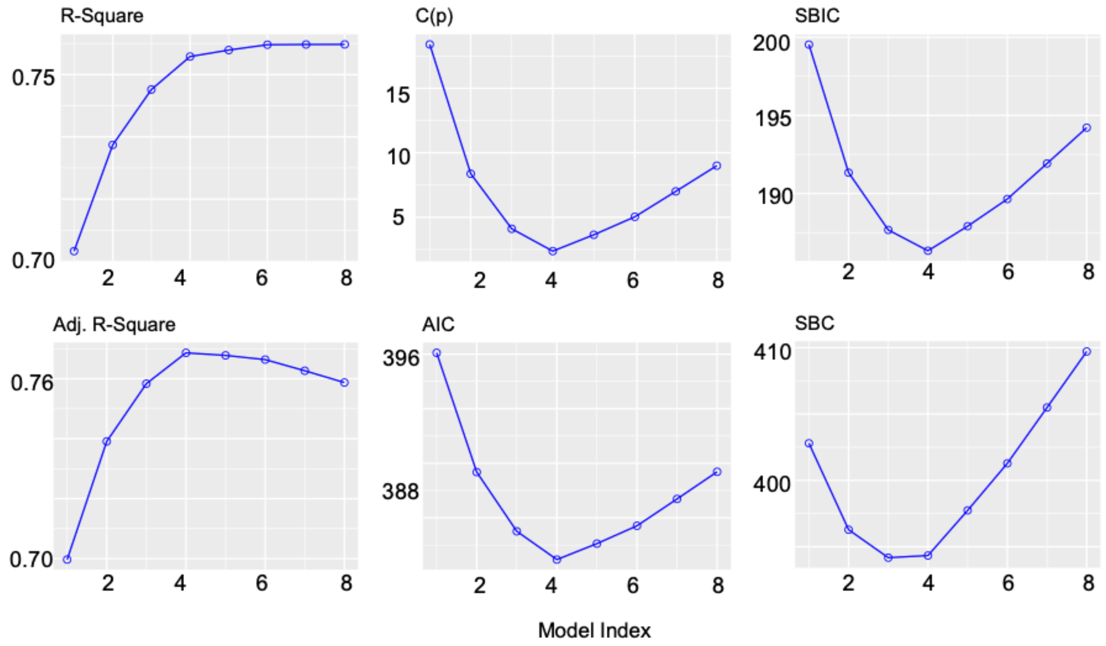


**Supplementary Figure 7.** Variable selection for the best-fitting linear model explaining the maximum variation in the rate of protein sequence evolution. **A**) Table containing the subset of predictors for each model tested. **B**) Various object criterion values for the models described in A.
